# Enzymatic synthesis of isotopically labeled hydrogen peroxide for mass spectrometry-based applications

**DOI:** 10.1101/2024.07.30.605865

**Authors:** Margaret Hoare, Ruiyue Tan, Isabella Militi, Kevin A. Welle, Kyle Swovick, Jennifer R. Hryhorenko, Sina Ghaemmaghami

## Abstract

Methionine oxidation is involved in multiple biological processes including protein misfolding and enzyme regulation. However, it is often challenging to measure levels of methionine oxidation by mass spectrometry, in part due to the prevalence of artifactual oxidation that occurs during sample preparation and ionization steps of typical proteomic workflows. Isotopically labeled hydrogen peroxide (H_2_^18^O_2_) can be used to block unoxidized methionines and enable accurate measurement of *in vivo* levels of methionine oxidation. However, H_2_^18^O_2_ is an expensive reagent that can be difficult to obtain from commercial sources. Here, we report a method for synthesizing H_2_^18^O_2_ in house. Glucose oxidase catalyzes the oxidation of β-D-glucose and produces hydrogen peroxide in the process. We took advantage of this reaction to enzymatically synthesize H_2_^18^O_2_ from ^18^O_2_ and assessed its concentration, purity, and utility in measuring methionine oxidation levels by mass spectrometry.

## Introduction

Side chains of methionines are susceptible to oxidation by reactive oxygen species (ROS) or monooxygenases.^1–6^ This post-translational modification converts the nonpolar methionine residues (Met) to polar methionine sulfoxide residues (MetO).^1–4^ MetO formation can induce protein misfolding and has been linked to a number of neurodegenerative disorders and pathological aging.^1–4,6^ Additionally, regulated methionine oxidation can modulate diverse cellular processes and signaling pathways. ^6^ Due to its involvement in protein damage and functional regulation, global quantification of methionine oxidation can provide important insights into a number of diverse biological processes.

It is challenging to measure levels of *in vivo* methionine oxidation by mass spectrometry in part because unoxidized methionines can become spontaneously oxidized during typical proteomic workflows. ^1–4^ Selective blocking of unoxidized methionines can prevent this artifactual oxidation and allow for more accurate quantitation of methionine oxidation. An example of such an approach is Methionine Oxidation by Blocking (MObB) ^1–4^In MObB, unoxidized methionines within denatured proteins are fully oxidized with isotopically labeled hydrogen peroxide (H_2_^18^O_2_) and blocked from spontaneous oxidation in subsequent steps of bottom-up proteomic workflows.^1,2^ Relative levels of ^16^O- and ^18^O-modified peptides can then be measured and used to determine levels of endogenously oxidized methionines.^1,2^ In addition to its utility in quantifying methionine oxidation, H_2_^18^O_2_ has been employed in other mass spectrometric applications including quantitation of H_2_O_2_-producing reactions, analysis of the effects of H_2_O_2_ on metabolite synthesis, and quantitation of H_2_O_2-_induced oxidation of macromolecules.^7–9^

Despite its usefulness in diverse mass spectrometric applications, H_2_^18^O_2_ is expensive and can be difficult to obtain from commercial sources. For example, the sale of H_2_^18^O_2_ was entirely discontinued between 2021 and 2023, and currently there is only a single supplier for this reagent (Sigma). Traditionally, hydrogen peroxide is synthesized through the hydrogenation and subsequent oxidation of anthraquinone, an aromatic organic compound that acts as a catalyst in this reaction.^10,11^ In the most popular synthetic methods, anthraquinone is hydrogenated by a trickle bed with a palladium catalyst. The hydrogenated anthraquinone is then oxidized by O_2_ through a bubble column, restoring anthraquinone and producing hydrogen peroxide is the process.^10^ The synthesized hydrogen peroxide is extracted using a sieve-plate extraction tower before it is purified via distillation.^10^ Alternative synthesis methods involving electrodes, biochemical approaches, or electrosynthesis reactions with different catalysts have also been reported.^11^

Enzymatic synthesis provides a more practical approach for in-house generation of H_2_^18^O_2_ in typical biochemical laboratories. Glucose oxidase (GOx), an oxidoreductase that originates from insects and fungi, has been used to generate H_2_O_2_ in multiple industries including pharmaceuticals, food, textiles, and biofuels.^12–16^ GOx catalyzes the oxidation of β-D-glucose to D-glucono-δ-lactone using a FAD cofactor and generates H_2_O_2_ by reduction of molecular oxygen (O_2_).^5,12–14,16–18^ Previously, GOx from *Aspergillus niger* has been used to generate ∼11 mM H_2_O_2_ for use in textile bleaching studies.^15^ In this study, we have optimized this enzymatic reaction and used ^18^O_2_ as substrate to generate ∼200 mM H_2_^18^O_2_ with high isotopic purity. We further demonstrated the efficacy of in-house generated H_2_^18^O_2_ in conducting quantitative mass spectrometric analyses of methionine oxidation levels.

## Results and Discussion

### Formation and characterization of H_2_^18^O_2_ generated by GOx

The mechanism for production of hydrogen peroxide by glucose oxidase is illustrated in Figure 1A. We utilized this enzymatic reaction to synthesize H_2_^18^O_2_ from ^18^O_2_ as described in detail in the Experimental Section (Figure 1B). In brief, the enzymatic conversion of β-D-glucose to D-glucono-δ-lactone was carried out in presence of a slow flow of ^18^O_2_ into an initially degassed solution containing GOx. In initial experiments we observed that the evolution of H_2_O_2_ deactivates GOx over time. Thus, to maximize the yield of H_2_O_2_, the reaction was supplemented with additional GOx after an initial generation period of 3.5 hours, allowing further generation of H_2_O_2_ for an additional hour. Following the reaction, GOx was removed from the mixture by filtration.

**Figure 1.**
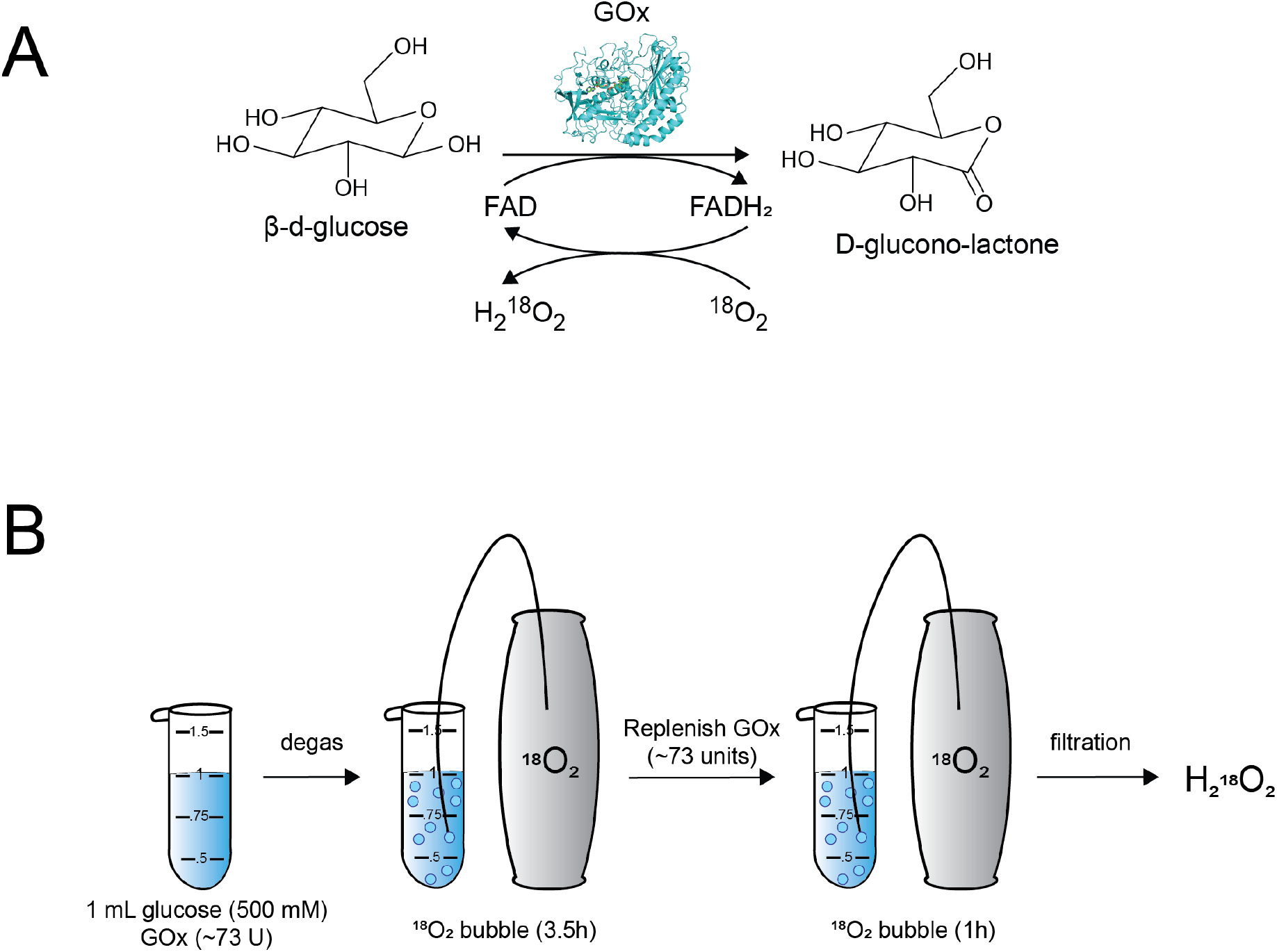
Reaction of glucose oxidase with β-D-glucose and ^18^O_2_ generates H_2_^18^O_2_. **A)** The reaction mechanism of glucose oxidase. **B)** Experimental protocol used for generation of H_2_^18^O_2_. Details of the protocol are described in the Experimental Section.

The purity and concentration of generated H_2_O_2_ was measured by mass spectrometry. An unoxidized synthetic peptide was fully oxidized with the synthesized H_2_^18^O_2_ or commercially obtained H_2_16O_2_ (Figure 2A). Relative levels of unmodified, ^16^O-modified and ^18^O-modified peptides were measured by mass spectrometry (Figure 2B). The peptide oxidized with in-house generated H_2_^18^O_2_ contained ∼94% ^18^O-labeled methionines, indicative of the isotopic purity of the oxidant. In a second experiment, the peptide was partially oxidized with known concentrations of H_2_^16^O_2_ and the resulting oxidation levels, as measured by mass spectrometry, were compared to peptides oxidized with various dilutions of in-house generated H_2_^18^O_2_. This comparison indicated that the in-house generated H_2_^18^O_2_ had a concentration of ∼230mM (Figure 2C).

**Figure 2.**
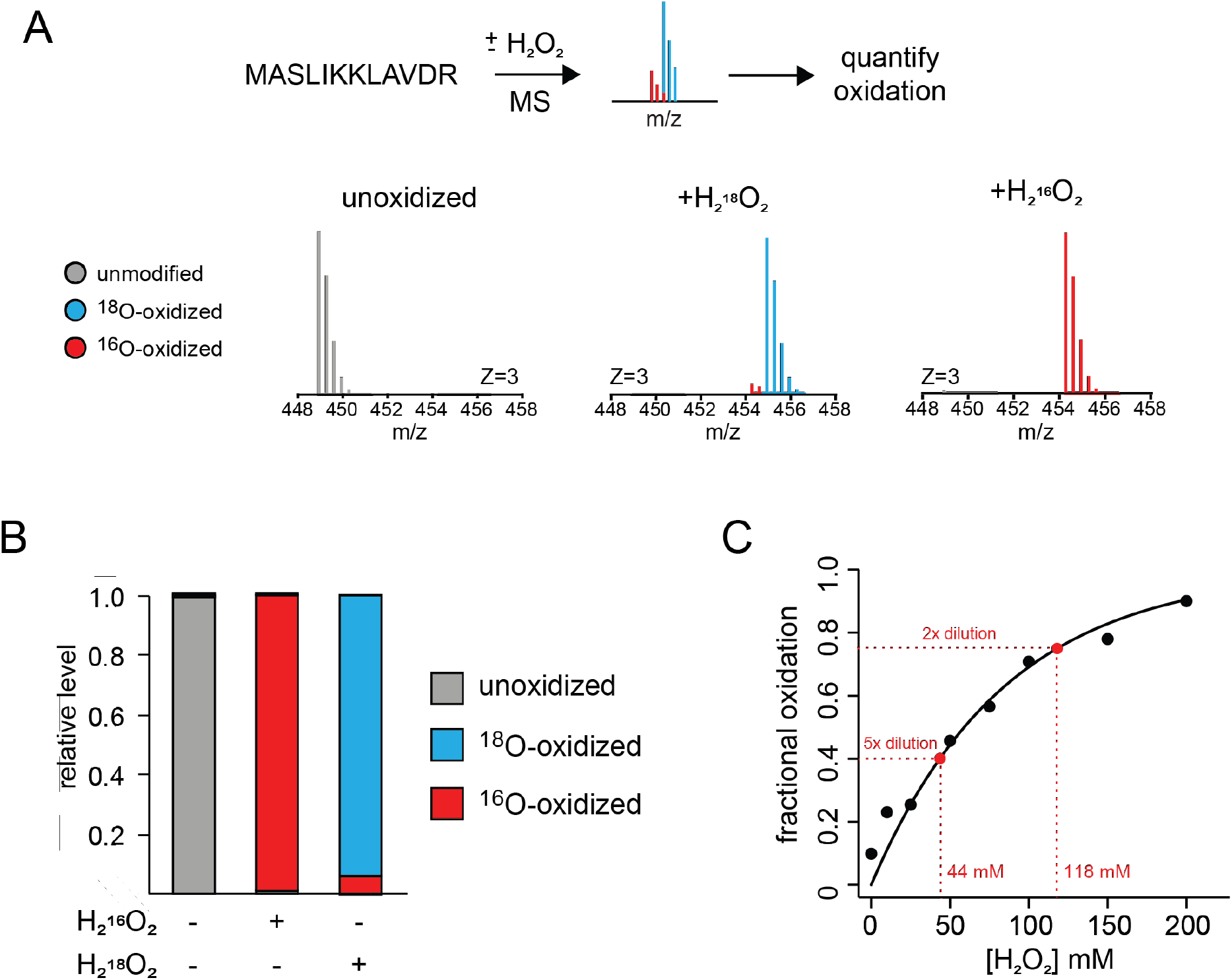
Synthesized H_2_^18^O_2_ has high isotopic purity and concentration. **A-B)** A synthetic peptide was oxidized with the in-house generated H_2_^18^O_2_ (A) and shown to be 94% ^18^O-labeled (B). **C)** The concentration of in-house generated H_2_^18^O_2_ was determined to be ∼230 mM by comparing its efficiency in oxidizing a model peptide with known concentrations of H_2_16O_2_ ladder.

### Generated H_2_^18^O_2_ can be used to accurately measure methionine oxidation levels

Next, we demonstrated that the generated H_2_^18^O_2_ can be used as an effective blocking agent, enabling the accurate measurement of methionine oxidation levels. A synthetic peptide was fully reduced by Methionine Sulfoxide Reductase A and Methionine Sulfoxide Reductase B (Msrs) or fully oxidized with H_2_16O_2._ Oxidized peptide was mixed with Msr-treated unoxidized peptide at variable ratios to generate mixtures with known pre-determined methionine oxidation levels. These mixtures were then oxidized with diluted H_2_^18^O_2_, resulting in ^18^O-oxidation of the previously unoxidized the methionines. Relative ^16^O-oxidation levels of each peptide mixture were determined by measuring the fractional populations of ^16^O- and ^18^O-oxidized peptides. The pairwise comparison of expected versus measured ^16^O-oxidation levels of peptide mixtures is shown in Figure 3. This analysis demonstrates that in-house generated H_2_^18^O_2_ can be effectively used as an isotopically labeled blocking reagent for accurate measurement of methionine oxidation levels.

**Figure 3.**
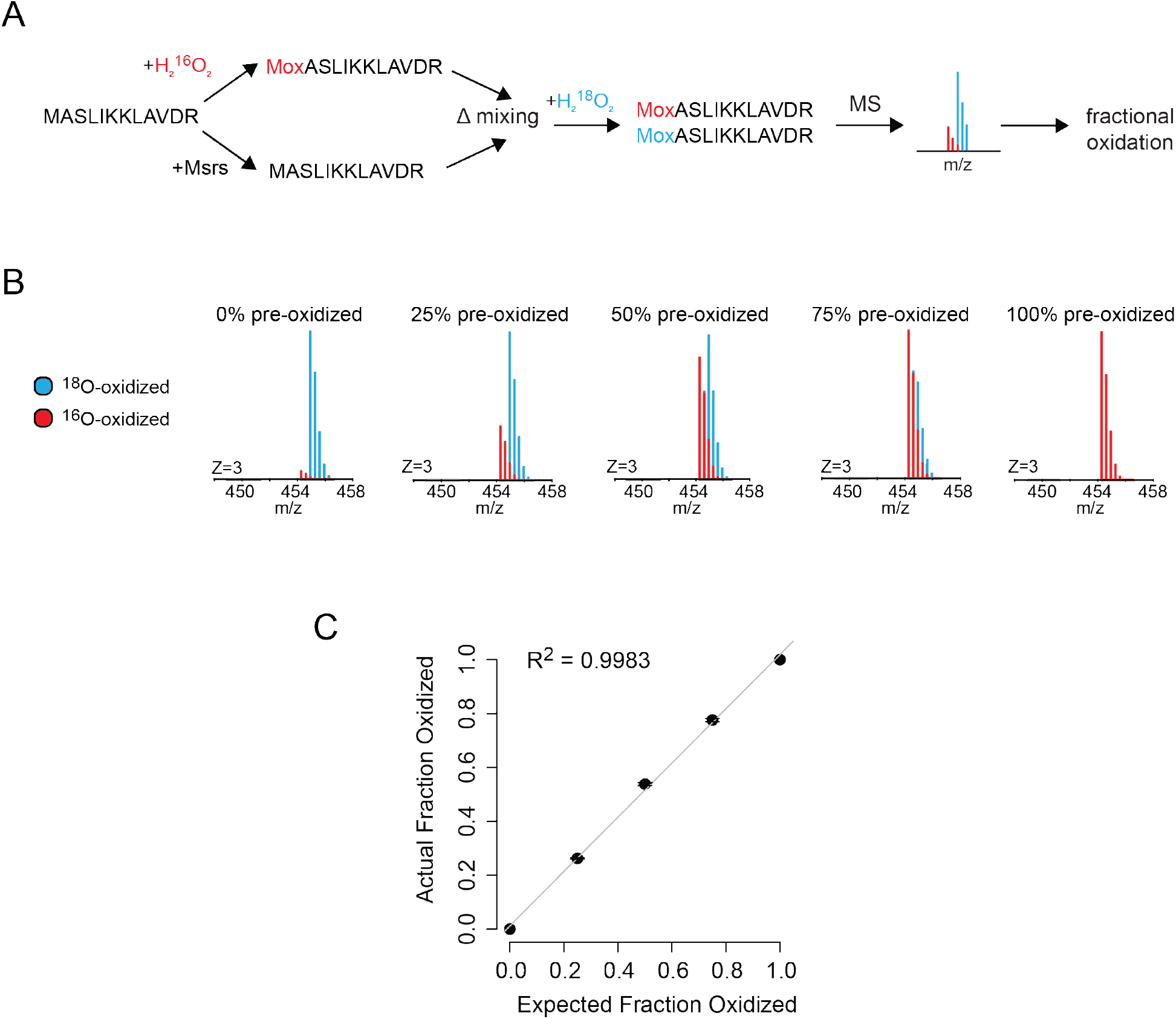
Synthesized H_2_^18^O_2_ can be used to accurately measure levels of methionine oxidation by mass spectrometry. **A-B)** Pre-oxidized peptide mixtures containing different levels of ^16^O-methionines (0%, 25%, 50%, 75%, 100%) were fully oxidized with H_2_^18^O_2_ **(A)** and analyzed by mass spectrometry **(B). C)** Fractional oxidation with ^16^O was measured and normalized to unoxidized and oxidized controls. The pairwise plot shows the correlation between measured and expected ^16^O-oxidation levels for each mixture. The error bars indicate standard deviations of two replicate experiments.

In summary, this study describes a protocol that employs a widely available enzyme, glucose oxidase, for generation of highly concentrated and isotopically enriched ^18^O-labeled hydrogen peroxide. The synthesized H_2_^18^O_2_ can be used to block unoxidized methionines and measure levels of methionine oxidation in mass spectrometric workflows such as MObB (Figure 3). However, although the described method for in-house synthesis of hydrogen peroxide is straightforward and accessible, there are two important caveats that require special consideration. First, the isotopic purity of the H_2_^18^O_2_ produced is dependent on the purity of the dissolved ^18^O_2_ in the reaction buffer. Thus, to obtain isotopically pure H_2_^18^O_2,_ removal of ^16^O_2_ by careful initial degassing, and subsequent use of highly pure ^18^O_2_ as a substrate is required. Second, the H_2_^18^O_2_ generated using the described protocol will also contain buffer components (in this case, sodium acetate) and glucose in oxidized and unoxidized forms. These impurities were inconsequential to the methionine blocking applications investigated in this study. However, if downstream applications require chemically pure H_2_^18^O_2_, further purification of the generated product may be required.

## Materials and methods

### Generation of hydrogen peroxide

1.5 mL of H_2_^18^O (Cambridge Isotope Laboratories, OLM-240-10G) was degassed in a 2 mL Eppendorf tube inside of a sealed vacuum flask connected to a vacuum. Glucose and sodium acetate were added to 1 mL of H_2_^18^O to attain final concentrations of 500 mM and 50 mM, respectively. 72.8 activity units of glucose oxidase (Sigma, G2133-10KU) were added to the solution, then ^18^O_2_ (Sigma, 602892-1L) was slowly bubbled from a pipette tip attached to tubing connected to a 1 L gas tank for 4.5 hours at 35°C. After 3.5 hours, another 72.8 units of glucose oxidase were added to the tube and the slow bubbling of ^18^O_2_ was continued for an additional hour. After incubation, the hydrogen peroxide was purified through centrifugation at 14,000xg for 20 minutes at 4 °C in a 0.5mL Amicon filter (3kDa MWCO) to remove the glucose oxidase. The synthesized H_2_^18^O_2_ was aliquoted and stored at -20 °C until use.

### Determination of the purity and concentration of H_2_^18^O_2_

To determine the purity of H_2_^18^O_2,_ 10 μg of a synthetic peptide (MASLIKKLAVDR) was incubated with diluted H_2_^18^O_2_ at 37 °C for 30 minutes and compared to an unoxidized control. Samples were frozen and lyophilized then desalted through homemade C18 columns and eluted in 50/50% acetonitrile (ACN)/H_2_O in 0.1% formic acid (FA). Samples were analyzed by direct injection mass spectrometry as described below. Raw MS data was analyzed using XCailbur (Thermo). Relative levels of unmodified, ^16^O- and ^18^O-oxidized peptides were determined by analyzing the intact mass spectra as described previously.^19^

To determine the concentration of H_2_^18^O_2,_ 10 μg of a synthetic peptide (MASLIKKLAVDR) was incubated with differing concentrations of H_2_^16^O_2_ (Fisher Bioreagents) or a 2x or 5x dilution of the generated H_2_^18^O_2_ solution for 10 minutes at 37 °C. Oxidation was quenched with 400 mM sodium sulfite and samples were frozen and lyophilized. Samples were desalted through homemade C18 columns and eluted in 50/50% ACN/H_2_O in 0.1% FA and analyzed by direct injection mass spectrometry as described below. Fractional oxidation was determined by least squares fitting the ^16^O-oxidation standard curve to a single exponential equation with the software KaleidaGraph and using the resulting fitted equation to determine the concentrations of the ^18^O-oxidized peptides.

### Generation of peptide mixtures with different oxidation levels

To generate fully oxidized peptides, 25 μg (372 μM) of MASLIKKLAVDR at a concentration of 0.5 μg/μl was oxidized with 160 mM H_2_O_2_ for 30 minutes at 37 °C. The sample was frozen and lyophilized to remove the hydrogen peroxide, then desalted in a homemade C18 column and eluted in 50% ACN/H_2_O in 0.1% trifluoroacetic acid (TFA). To generate fully unoxidized peptides, 25 μg of the peptide was incubated with 50 mM dithiothreitol (Sigma), 1.25 μM of Methionine Sulfoxide Reductase A and 12.3 μM of Methionine Sulfoxide Reductase B that were recombinantly expressed and purified from *E*.*coli* ^20,21^ in 50 mM Tris buffer for 45 minutes at 37 °C. Samples were lyophilized then desalted to remove enzymes and salts. After desalting, fully oxidized and fully reduced peptides were mixed to generate specific fractional oxidations. These pre-oxidized mixtures were then frozen and lyophilized before reconstitution in 1.25x diluted H_2_^18^O_2_. The peptides were oxidized for 30 minutes at 37 °C, then frozen and lyophilized before desalting with homemade C18 columns into 50/50% ACN/H_2_O in 0.1% FA. The sample was diluted to 15 μg/mL before analysis by mass spectrometry.

### Mass Spectrometry

Peptides were diluted in 50/50% ACN/H_2_O in 0.1% FA to 15 μg/mL. 40 μl of this peptide was directly injected into a Q Exactive Plus Mass Spectrometer (Thermo Fisher) using a Dionex Ultimate 3000 HPLC (Thermo Fisher) with a flow rate of 100 μL/min. The solvent was a 50/50% mixture of 0.1% FA in H_2_O and 0.1% FA in ACN that was injected for 3 minutes. A HESI source set in positive mode ionized the peptides at resolution of 70,000 at m/z 200 with a 240 ms maximum injection time, AGC target of 1e6, and an overall range of 300-2000 m/z. MS1 files were generated from raw files using MSConvert^22^ and fractional oxidations were measured as previously described.^19^ We note that under some conditions, a small population of the peptide was oxidized to form methionine sulfones in addition to methionine sulfoxides. In these cases, fractional oxidations were measured using the population averaged ratios of ^16^O/^18^O-oxidized relative to ^18^O/^18^O-oxidized peptides, as well as ^16^O-oxidized relative to ^18^O-oxidized peptides.

## Author Information

### Author Contributions

The study concept was conceived by M.H., R.T. and S.G. The experiments were carried out by M.H., I.M., and R.T. Mass spectrometry was performed by K.W., K.S. and J.H. Data analysis was conducted by M.H., R.T., and S.G. The initial draft of the manuscript was written by M.H. and S.G.

## Abbreviations

ROS: reactive oxygen species
Msrs: methionine sulfoxide reductases
MObB: Methionine Oxidation by Blocking
GOx: glucose oxidase
FAD: flavin adenine dinucleotide
H_2_O_2_: hydrogen peroxide
Met: methionine
MetO: methionine sulfoxide
ACN: acetonitrile
FA: formic acid
TFA: trifluoroacetic acid
HESI: heated electrospray ionization
AGC: automatic gain control

## References

(1) Bettinger, J. Q.; Welle, K. A.; Hryhorenko, J. R.; Ghaemmaghami, S. Quantitative Analysis of in Vivo Methionine Oxidation of the Human Proteome. J. Proteome Res. 2020, 19 (2), 624–633. 10.1021/acs.jproteome.9b00505.

(2) Bettinger, J. Q.; Simon, M.; Korotkov, A.; Welle, K. A.; Hryhorenko, J. R.; Seluanov, A.; Gorbunova, V.; Ghaemmaghami, S. Accurate Proteomewide Measurement of Methionine Oxidation in Aging Mouse Brains. J. Proteome Res. 2022, 21 (6), 1495–1509. 10.1021/acs.jproteome.2c00127.

(3) Liu, H.; Ponniah, G.; Neill, A.; Patel, R.; Andrien, B. Accurate Determination of Protein Methionine Oxidation by Stable Isotope Labeling and LC-MS Analysis. Anal. Chem. 2013, 85 (24), 11705–11709. 10.1021/ac403072w.

(4) Shipman, J. T.; Go, E. P.; Desaire, H. Method for Quantifying Oxidized Methionines and Application to HIV-1 Env. J. Am. Soc. Mass Spectrom. 2018, 29 (10), 2041–2047. 10.1007/s13361-018-2010-2.

(5) Kleppe, K. The Effect of Hydrogen Peroxide on Glucose Oxidase from Aspergillus Niger *. Biochemistry 1966, 5 (1), 139–143. 10.1021/bi00865a018.

(6) Hoshi, T.; Heinemann, S. H. Regulation of Cell Function by Methionine Oxidation and Reduction. J Physiol 2001, 531 (Pt 1), 1–11. 10.1111/j.1469-7793.2001.0001j.x.

(7) Liu, X.; Cooper, D. E.; Cluntun, A. A.; Warmoes, M. O.; Zhao, S.; Reid, M. A.; Liu, J.; Lund, P. J.; Lopes, M.; Garcia, B. A.; Wellen, K. E.; Kirsch, D. G.; Locasale, J. W. Acetate Production from Glucose and Coupling to Mitochondrial Metabolism in Mammals. Cell 2018, 175 (2), 502-513.e13. 10.1016/j.cell.2018.08.040.

(8) Vermilyea, A. W.; Dixon, T. C.; Voelker, B. M. Use of H2(18)O2 to Measure Absolute Rates of Dark H2O2 Production in Freshwater Systems. Environ Sci Technol 2010, 44 (8), 3066–3072. 10.1021/es100209h.

(9) Hofer, T.; Badouard, C.; Bajak, E.; Ravanat, J.-L.; Mattsson, Å.; Cotgreave, I. A. Hydrogen Peroxide Causes Greater Oxidation in Cellular RNA than in DNA. 2005, 386 (4), 333–337. 10.1515/BC.2005.040.

(10) Chen, Q. Development of an Anthraquinone Process for the Production of Hydrogen Peroxide in a Trickle Bed Reactor—From Bench Scale to Industrial Scale. Chemical Engineering and Processing: Process Intensification 2008, 47 (5), 787–792. 10.1016/j.cep.2006.12.012.

(11) Perry, S. C.; Pangotra, D.; Vieira, L.; Csepei, L.-I.; Sieber, V.; Wang, L.; Ponce de León, C.; Walsh, F. C. Electrochemical Synthesis of Hydrogen Peroxide from Water and Oxygen. Nat Rev Chem 2019, 3 (7), 442–458. 10.1038/s41570-019-0110-6.

(12) Bauer, J. A.; Zámocká, M.; Majtán, J.; Bauerová-Hlinková, V. Glucose Oxidase, an Enzyme “Ferrari”: Its Structure, Function, Production and Properties in the Light of Various Industrial and Biotechnological Applications. Biomolecules 2022, 12 (3), 472. 10.3390/biom12030472.

(13) Bankar, S. B.; Bule, M. V.; Singhal, R. S.; Ananthanarayan, L. Glucose Oxidase — An Overview. Biotechnology Advances 2009, 27 (4), 489–501. 10.1016/j.biotechadv.2009.04.003.

(14) Khatami, S. H.; Vakili, O.; Ahmadi, N.; Soltani Fard, E.; Mousavi, P.; Khalvati, B.; Maleksabet, A.; Savardashtaki, A.; Taheri-Anganeh, M.; Movahedpour, A. Glucose Oxidase: Applications, Sources, and Recombinant Production. Biotechnology and Applied Biochemistry 2022, 69 (3), 939–950. 10.1002/bab.2165.

(15) Tzanov, T.; Costa, S. A.; Gübitz, G. M.; Cavaco-Paulo, A. Hydrogen Peroxide Generation with Immobilized Glucose Oxidase for Textile Bleaching. Journal of Biotechnology 2002, 93 (1), 87–94. 10.1016/S0168-1656(01)00386-8.

(16) Dubey, M. K.; Zehra, A.; Aamir, M.; Meena, M.; Ahirwal, L.; Singh, S.; Shukla, S.; Upadhyay, R. S.; Bueno-Mari, R.; Bajpai, V. K. Improvement Strategies, Cost Effective Production, and Potential Applications of Fungal Glucose Oxidase (GOD): Current Updates. Front Microbiol 2017, 8, 1032. 10.3389/fmicb.2017.01032.

(17) Bao, J.; Furumoto, K.; Yoshimoto, M.; Fukunaga, K.; Nakao, K. Competitive Inhibition by Hydrogen Peroxide Produced in Glucose Oxidation Catalyzed by Glucose Oxidase. Biochemical Engineering Journal 2003, 13 (1), 69–72. 10.1016/S1369-703X(02)00120-1.

(18) Hecht, H. J.; Kalisz, H. M.; Hendle, J.; Schmid, R. D.; Schomburg, D. Crystal Structure of Glucose Oxidase from Aspergillus Niger Refined at 2·3 Å Reslution. Journal of Molecular Biology 1993, 229 (1), 153–172. 10.1006/jmbi.1993.1015.

(19) Tan, R.; Hoare, M.; Welle, K. A.; Swovick, K.; Hryhorenko, J. R.; Ghaemmaghami, S. Folding Stabilities of Ribosome-Bound Nascent Polypeptides Probed by Mass Spectrometry. Proceedings of the National Academy of Sciences 2023, 120 (33), e2303167120. 10.1073/pnas.2303167120.

(20) Tan, R.; Hoare, M.; Bellomio, P.; Broas, S.; Camacho, K.; Swovick, K.; Welle, K. A.; Hryhorenko, J. R.; Ghaemmaghami, S. Formylation Facilitates the Reduction of Oxidized Initiator Methionines. bioRxiv February 7, 2024, p 2024.02.06.579201. 10.1101/2024.02.06.579201.

(21) Hoare, M.; Tan, R.; Welle, K. A.; Swovick, K.; Hryhorenko, J. R.; Ghaemmaghami, S. Methionine Alkylation as an Approach to Quantify Methionine Oxidation Using Mass Spectrometry. J Am Soc Mass Spectrom 2024, 35 (3), 433–440. 10.1021/jasms.3c00337.

(22) Chambers, M. C.; Maclean, B.; Burke, R.; Amodei, D.; Ruderman, D. L.; Neumann, S.; Gatto, L.; Fischer, B.; Pratt, B.; Egertson, J.; Hoff, K.; Kessner, D.; Tasman, N.; Shulman, N.; Frewen, B.; Baker, T. A.; Brusniak, M.-Y.; Paulse, C.; Creasy, D.; Flashner, L.; Kani, K.; Moulding, C.; Seymour, S. L.; Nuwaysir, L. M.; Lefebvre, B.; Kuhlmann, F.; Roark, J.; Rainer, P.; Detlev, S.; Hemenway, T.; Huhmer, A.; Langridge, J.; Connolly, B.; Chadick, T.; Holly, K.; Eckels, J.; Deutsch, E. W.; Moritz, R. L.; Katz, J. E.; Agus, D. B.; MacCoss, M.; Tabb, D. L.; Mallick, P. A Cross-Platform Toolkit for Mass Spectrometry and Proteomics. Nat Biotechnol 2012, 30 (10), 918–920. 10.1038/nbt.2377.

